# Mismatch repair MLH complexes make distinct contributions to post-replicative mismatch repair versus trinucleotide repeat expansions

**DOI:** 10.64898/2026.01.20.700715

**Authors:** Katherine M. Casazza, Greg M. Williams, Lauren Johengen, Maxwell Keller, Luke D. Hess, Samantha Phelps, Natalie A. Lamb, Jennifer A. Surtees

## Abstract

Mismatch repair (MMR) is a highly conserved DNA repair pathway that promotes genome stability by directing the repair of errors in DNA replication. In *Saccharomyces cerevisiae*, MMR is initiated by either Msh2-Msh3 or Msh2-Msh6, via recognition of insertion deletion loops (IDLs; up to ∼ 17 nucleotides) and misincorporation events, respectively. Both complexes recognize and bind small (1-2 nucleotide) IDLs. Once bound, MSH complexes recruit one or more downstream MLH complexes to continue repair: Mlh1-Pms1, Mlh1-Mlh2 and/or Mlh1-Mlh3. Msh2-Msh3 also promotes *CAG* trinucleotide repeat (TNR) expansions through specific DNA-binding to TNR DNA structures, followed by recruitment of MLH complexes. These expansions lead to genome instability that causes neurodegenerative diseases such as Huntington’s Disease in humans. Here, we defined a hierarchy of MLH function in these Msh2-Msh3-mediated pathways *in vivo* in *S. cerevisiae*. We determined that Mlh1-Pms1 is the primary MLH complex required in Msh2-Msh3-mediated MMR. In contrast, all three MLH complexes were required to promote *CAG* expansions, with loss of Mlh1-Pms1 or Mlh1-Mlh2 exhibiting the strongest effects. Mutations in *PMS1* and *MLH3* were synergistic. We propose a model in which Mlh1-Pms1 is primarily responsible for “appropriate” Msh2-Msh3-mediated MMR, while all three MLH complexes collaborate specifically in the presence of *CAG* structure, to promote a “pathogenic” Msh2-Msh3-mediated pathway that leads to expansions. Our model highlights the importance of DNA structure-dependent conformations in modulating MLH function.

## Introduction

Mismatch repair (MMR) functions at the replication fork to correct errors in DNA replication, thus preventing their incorporation into the genome, minimizing mutagenesis and protecting higher eukaryotic cells from carcinogenesis (Kunkel and Erie 2005; Jiricny 2006; Reyes *et al*. 2015). In most eukaryotes, MMR is initiated by one of two MutS homolog (MSH) heterodimers, Msh2-Msh6 and Msh2-Msh3. These complexes have separate but overlapping substrate specificity for distinct DNA structures; Msh2-Msh6 recognizes and binds nucleotide misincorporations and small insertion/deletion loops (IDLs), while Msh2-Msh3 recognizes and binds IDLs up to ∼17 nucleotides, as well as a subset of base-base mispairs (Strand *et al*. 1995; Marsischky *et al*. 1996; Kirkpatrick and Petes 1997; Flores-Rozas and Kolodner 1998; Genschel *et al*. 1998; Harfe and Jinks-Robertson 1999; Gragg *et al*. 2002). With this broad specificity, the mismatch repair system is able to recognize and direct repair of a variety of replication errors (Sia *et al*. 1997; Kunkel and Erie 2005; Lee *et al*. 2007; Warren *et al*. 2007; Gupta *et al*. 2011). After error recognition and binding, MSH complexes recruit MutL homolog (MLH) heterodimers. In most eukaryotes, including *Saccharomyces cerevisiae*, there are three MLH complexes that contribute to MMR. In yeast, the Mlh1 subunit forms complexes with Pms1 (Pms2 in mammals), Mlh2 (Pms1 in mammals) and Mlh3 to form MutLα, MutLβ and MutLγ, respectively (Kunkel and Erie 2005; Pannafino and Alani 2021). Mlh1-Pms1 and Mlh1-Mlh3 each have latent endonuclease activity that is stimulated upon recruitment to a mispair by Msh2-Msh6 or Msh2-Msh3 and by interactions with PCNA. MLH endonuclease activity results in nicking that is directed to the nascent strand to remove the error and resynthesize with DNA polymerase δ (Furman *et al*. 2021; Pannafino and Alani 2021). *In vivo* reporter assays indicate that Mlh1-Pms1 (MutLα) plays a critical role in both Msh2-Msh3 and Msh2-Msh6-mediated MMR (Prolla *et al*. 1994; Habraken *et al*. 1997; Kadyrov *et al*. 2007), although much of this work has been done with reporter substrates whose repair can be directed by either Msh2-Msh6 or Msh2-Msh3. Mlh1-Mlh3 and Mlh1-Mlh2 also contribute to MMR but play less prominent roles; their loss leads to only small decreases in MMR. Notably, the role of Mlh1-Mlh2 or Mlh1-Mlh3 has been primarily evaluated in Msh2-Msh6-mediated repair; MLH function in Msh2-Msh3-mediated MMR has been less well-characterized, Although Mlh1-Mlh3 has been implicated primarily in Msh2-Msh3-mediated MMR (Flores-Rozas and Kolodner 1998; Harfe *et al*. 2000; Lipkin *et al*. 2000; Romanova and Crouse 2013; Al-Sweel *et al*. 2017). Genetic evidence suggested that Mlh1-Mlh3 may collaborate with Mlh1-Pms1 (Romanova and Crouse 2013), although the context for this cooperation remained unclear. We demonstrated that Msh2-Msh3 interacts with Mlh1-Mlh3 and stimulates its endonuclease activity (Rogacheva *et al*. 2014; Al-Sweel *et al*. 2017). Nonetheless, Mlh1-Mlh3’s primary function is in meiotic recombination, where it regulates the crossing-over of homologous chromosomes in an endonuclease-dependent manner (Lipkin *et al*. 2002; Nishant *et al*. 2008). Mlh1-Mlh2 does not have endonuclease activity and is predicted to act as an accessory factor in MMR (Harfe *et al*. 2000; Campbell *et al*. 2014). The mechanism for its activity remains uncharacterized, although Mlh1-Mlh2 interacted with both Msh2-Msh3 and Msh2-Msh6 *in vitro* (Campbell *et al*. 2014). Mlh1-Mlh2 also plays a prominent role in regulating gene conversions tract lengths in meiosis, through interactions with Mer3 and Pif1 (Duroc *et al*. 2017; Vernekar *et al*. 2021; Altmannova *et al*. 2023).

Defects in the MMR system are detrimental, increasing mutagenesis and leading to a strong predisposition to several cancers (Miao *et al*. 2015; DE’ Angelis *et al*. 2018; Santos *et al*. 2018). Thus, it is striking that components of the MMR system contribute to genomic instability in the context of trinucleotide repeat (TNR) expansions (Schmidt and Pearson 2016; Khristich *et al*. 2020; Lahue 2020; Iyer and Pluciennik 2021; Richard 2021), which are the underlying cause of over 40 neurodegenerative and neuromuscular disorders. The pathway to disease-causing expansions is thought to begin with incremental expansions of a normal triplet repeat tract to a threshold length that is more susceptible to further expansions (Higham *et al*. 2012; Du *et al*. 2013; Higham and Monckton 2013; Williams and Surtees 2015). Threshold length tracts then undergo further expansion to pathogenic lengths (Leeflang *et al*. 1995; Leeflang *et al*. 1999); threshold and pathogenic lengths are locus-dependent. Msh2-Msh3, but not Msh2-Msh6, contributes to *CAG/CTG* expansions in a variety of model and clinical systems (Owen *et al*. 2005; Foiry *et al*. 2006; Kantartzis *et al*. 2012; Keogh *et al*. 2017). In this study, we focused on *CAG* repeat expansions, which underlie Huntington’s disease (HD), spinocerebellar ataxias (SCAs) and other disorders (Paulson and Fischbeck 1996; Mirkin 2007; Yamada *et al*. 2008; Kacher *et al*. 2024). In transgenic mice carrying an expanded, pathogenic allele of the human Huntingtin gene (>100 *CAG* repeats in exon 1) (Mangiarini *et al*. 1996), inactivation of *Msh2* or *Msh3* reduced tract instability and eliminated further expansions (Manley *et al*. 1999; Owen *et al*. 2005). Also, in transgenic mice, SNPs identified in *Msh3* that increased *Msh3* expression correlated with increased tract instability (TOMÉ *et al*. 2013). In human cell lines, *Msh3* promoted expansions of *CAG* tracts and its expression level was correlated with *CAG* expansion rates (Gannon *et al*. 2012) (Keogh *et al*. 2017). These findings are consistent with GWAS studies that identified *Msh2* and *Msh3* as genetic modifiers of HD, demonstrating the importance of MMR to clinical disease (Consortium 2015; Lee *et al*. 2025). In *Saccharomyces cerevisiae*, we demonstrated that *msh3Δ* reduces *CAG* expansion rate, using a reporter that measures expansions in smaller, threshold-length tracts. (Kantartzis *et al*. 2012) and demonstrated a mechanism of incremental expansions at these tract-lengths (Williams and Surtees 2015). Notably, we previously demonstrated that the specific DNA mispair binding domain (MBD) of Msh3 was insufficient to promote TNR expansions, highlighting the importance of other components of MMR for this pathogenic process, including potentially interactions with MLH complexes (Casazza *et al*. 2025).

Consistent with this suggestion, several mammalian studies using large, expanded *CAG* tracts support the hypothesis that MLH complexes also contribute to expansions; this is predicted to be downstream of Msh2-Msh3 function. *Mlh1* was genetically linked to HD onset through GWAS studies and was required for somatic expansions in HD transgenic mice (Pinto *et al*. 2013; Consortium 2019). Sequestration of Mlh1 by FAN1 reduced rates of TNR expansions (Goold *et al*. 2021; Porro *et al*. 2021; Senoussi *et al*. 2025). Loss of mammalian Pms2 (Pms1 in yeast), Pms1 (Mlh2 in yeast) and Mlh3 reduced *CAG* expansions of pathogenic tracts in human cells and HD mouse models (Pinto *et al*. 2013; Hayward *et al*. 2024). Mlh1-Mlh3’s role in promoting expansions is dependent on its endonuclease activity (Roy *et al*. 2021). The requirement of all MLH complexes for promoting TNR expansions indicates a striking distinction between post-replicative Msh2-Msh3-medidated MMR pathway, which promotes genome stability, and Msh2-Msh3-mediated TNR expansions, i.e. genome instability. Here we compare directly the contributions of each MLH complex to Msh2-Msh3-mediated *CAG* repeat expansion, using threshold length tracts, and canonical post-replicative MMR pathways in *S. cerevisiae* and define a distinct hierarchy of MLH function in each Msh2-Msh3-mediated pathway.

## Materials and Methods

### Strains and Media

All plasmids and strains used in this study are described in Tables 1 and 2. All yeast transformations were performed using the lithium acetate method (Gietz *et al*. 1992). Yeast strains were derived from the S288c background; FY23 was used for slippage assays and FY86 was used for TNR expansion assays (Winston *et al*. 1995). *pms1Δ, mlh3Δ, mlh2Δ, mlh1Δ* and strains containing either mutation were constructed by amplifying a chromosomal mutant::*KANMX* fragment from the yeast deletion collection that was integrated into the respective chromosomal location in FY86 and FY23. *pms1E707K* and *mlh3D523N* mutants were created through Q5 site directed mutagenesis to generate integration plasmids pGW20 (*pms1E707K::NATMX*) and pYH5 (*mlh3D523N::KANMX)*. Sequences of both plasmids were confirmed through Sanger sequencing.

**Table 1:**
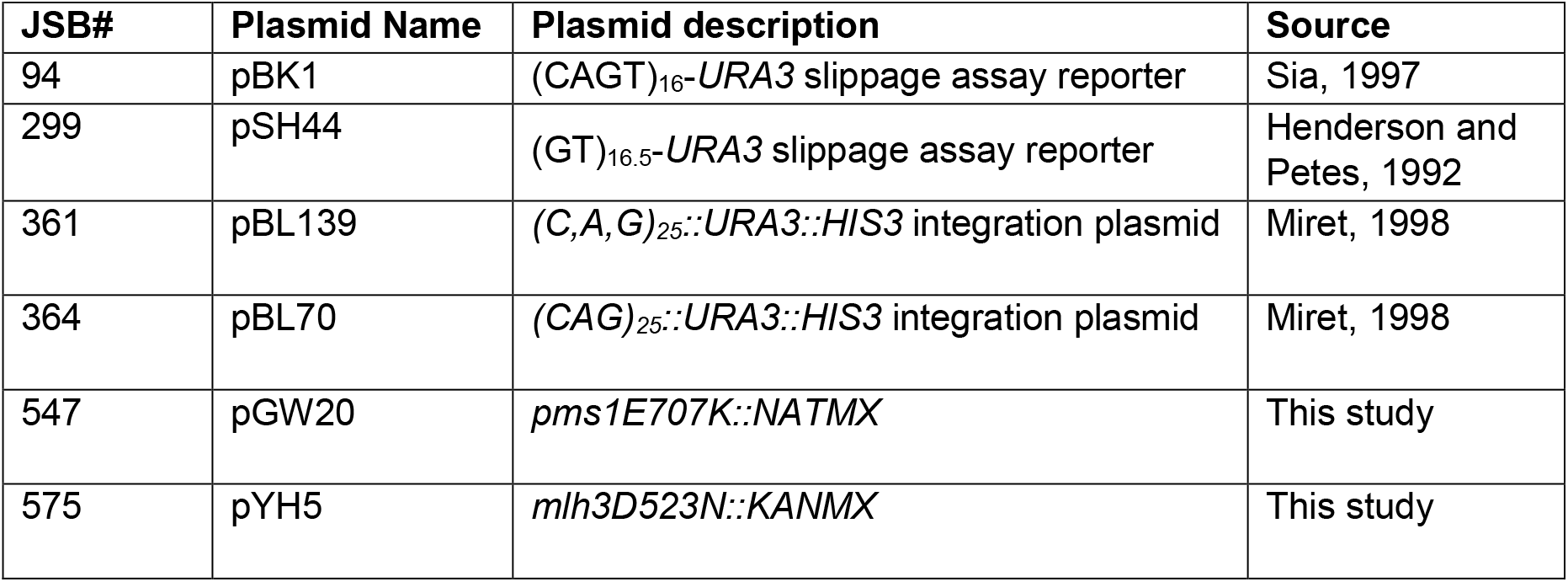
Plasmids used in this study.

| **Table 1: Plasmids used in this study** |  |  |  |
| --- | --- | --- | --- |
| **JSB#** | **Plasmid Name** | **Plasmid description** | **Source** |
| 94 | pBK1 | (CAGT)<sub>16</sub>-*URA3* slippage assay reporter | Sia, 1997 |
| 299 | pSH44 | (GT)<sub>16.5</sub>-*URA3* slippage assay reporter | Henderson and Petes, 1992 |
| 361 | pBL139 | (*C,A,G*)<sub>25</sub>::*URA3::HIS3* integration plasmid | Miret, 1998 |
| 364 | pBL70 | (*CAG*)<sub>25</sub>::*URA3::HIS3* integration plasmid | Miret, 1998 |
| 547 | pGW20 | *pms1E707K::NATMX* | This study |
| 575 | pYH5 | *mlh3D523N::KANMX* | This study |

**Table 2:**
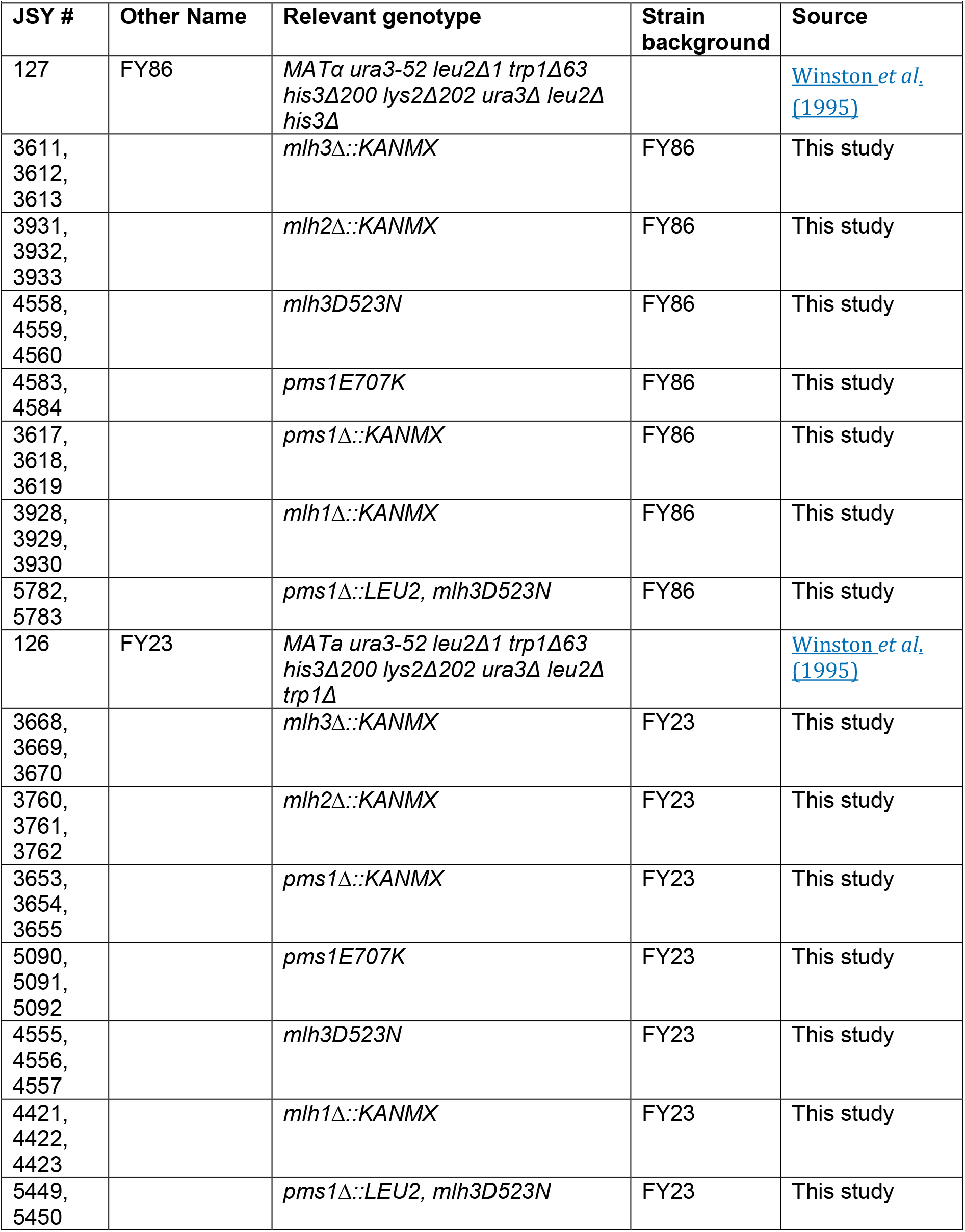
Yeast strains used in this study.

TNR substrates were integrated into all strains as described previously (Dixon *et al*. 2004; Williams and Surtees 2018). Two different substrates were used and integrated into the genetic backgrounds, with the following sequence on the lagging daughter strand: 1) a *(CAG)*_*25*_ repeat (pBL70) and 2) a scrambled *(C,A,G)*_*25*_ repeat (pBL139) (Dixon *et al*. 2004). Each repeat tract was cloned into the regulatory region controlling expression of the *URA3* reporter gene. When the distance between the TATA box and the initiator *ATG* for the *URA3* gene is increased beyond 29 repeats, *URA3* is no longer expressed, making the cells resistant to 5-FOA.

The microsatellite instability construct has been described previously(Sia *et al*. 1997). Repeating base units of 2n, consisting of *(GT*)_16.5_ or 4n, consisting of (*CAGT*)_16.5_ were placed upstream of *URA3*. At these lengths, *URA3* is in frame and cells are Ura^+^. If slippage events occur and are not repaired, the resulting insertion/deletion event shifts *URA3* out of frame. The resulting event makes the cells Ura^-^ and resistant to 5-FOA.

### Assay for TNR Expansion

TNR expansion assays were performed as described (WILLIAMS AND SURTEES). Briefly, single colonies were obtained on synthetic medium (SC) lacking histidine (SC-his) or lacking histidine and uracil (SC-his-ura) for *pms1Δ, pms1E707K, pms1Δ mlh3D523N*, and *mlh1Δ*. Individual colonies were selected from ≥3 independent isolates of each genetic background were assayed. Colonies with unexpanded tracts, which were confirmed by PCR analysis, were diluted and plated on SC-his and incubated at 30^°^C for 3-4 days to allow expansions to occur. Several ∼2mm colonies were selected, diluted and plated onto permissive (SC-his) and selective (SC-his +5-FOA) and incubated at 30^°^C from 3-4 days. Colonies were counted and expansion rates calculated as described (DRAKE). The 95% confidence intervals were determined from tables of confidence intervals for the median (NAIR 1940; DIXON AND MASSEY 1969). p-values were determined by Mann-Whitney rank analysis (NAIR 1940; DIXON AND MASSEY 1969; Drake 1991; Sia *et al*. 1997) in GraphPad Prism.

True expansions were determined as previously described (Williams and Surtees 2015; Williams and Surtees 2018; Williams *et al*. 2020), by amplifying the reporter promoter region of 5-FOA-resistant colonies with SO295 (AAACTCGGTTTGACGCCTCCCATG) and SO296 (AGCAACAGGACTAGGATGAGTAGC) and digesting with *SphI* to release the TNR tract. Tract mobility was assessed by electrophoresis through a 12% native polyacrylamide gel (0.5 X TBE).

### Microsatellite instability assay

Microsatellite instability assays were performed as described (Sia *et al*. 1997). Single colonies were grown on SC-tryptophan (SC-trp) to maintain the reporter plasmids. Colonies of ∼2 mm were selected from ≥3 independent isolates of each genetic background. Individual colonies were resuspended in 3 mL of liquid SC-trp and incubated with shaking for 20 hours at 30^°^C. Overnight cultures were serial diluted and plated on permissive (SC-trp) and selective (SC-trp +5-FOA) plates. Plates were incubated at 30ºC for 2-4 days. Mutation rates were calculated by method of the median (Drake 1991). 95% confidence intervals were determined using tables of confidence intervals for the median (NAIR 1940; DIXON AND MASSEY 1969). p-values were determined by Mann-Whitney rank analysis in GraphPad Prism.

## Results

### MLH complexes make distinct contributions to Msh2-Msh3-mediated MMR

The roles of MLH complexes in MMR have largely been tested in reporter assays that measure correction of overlapping Msh2-Msh3 and Msh2-Msh6 repair substrates (Sia *et al*. 1997; Flores-Rozas and Kolodner 1998; Romanova and Crouse 2013; Campbell *et al*. 2014; Rogacheva *et al*. 2014). In this study, we were specifically concerned with defining the requirement for each of the three MLH complexes in Msh2-Msh3-mediated MMR. Therefore, we used a set of reporter plasmids that measures frameshift or “slippage” events in the presence of dinucleotide (*GT)*_*16*.*5*_ (DAI *et al*.) or tetranucleotide (*CAGT*)_16.5_ repeat plasmids (Sia *et al*. 1997) because *MSH3* plays a critical role in repairing IDLs resulting from repeats of these sizes (Sia *et al*. 1997; Lee *et al*. 2007; Kumar *et al*. 2013). Further, slippage events occur with these size loops at high enough rates that a range of changes can potentially be observed and compared between genotypes **(Figure 1**). The effects of deletion of *MLH1* and *PMS1* on dinucleotide slippage rates have been previously tested, but tetranucleotide repeats have not been (Hall *et al*. 2001).

**Figure 1.**
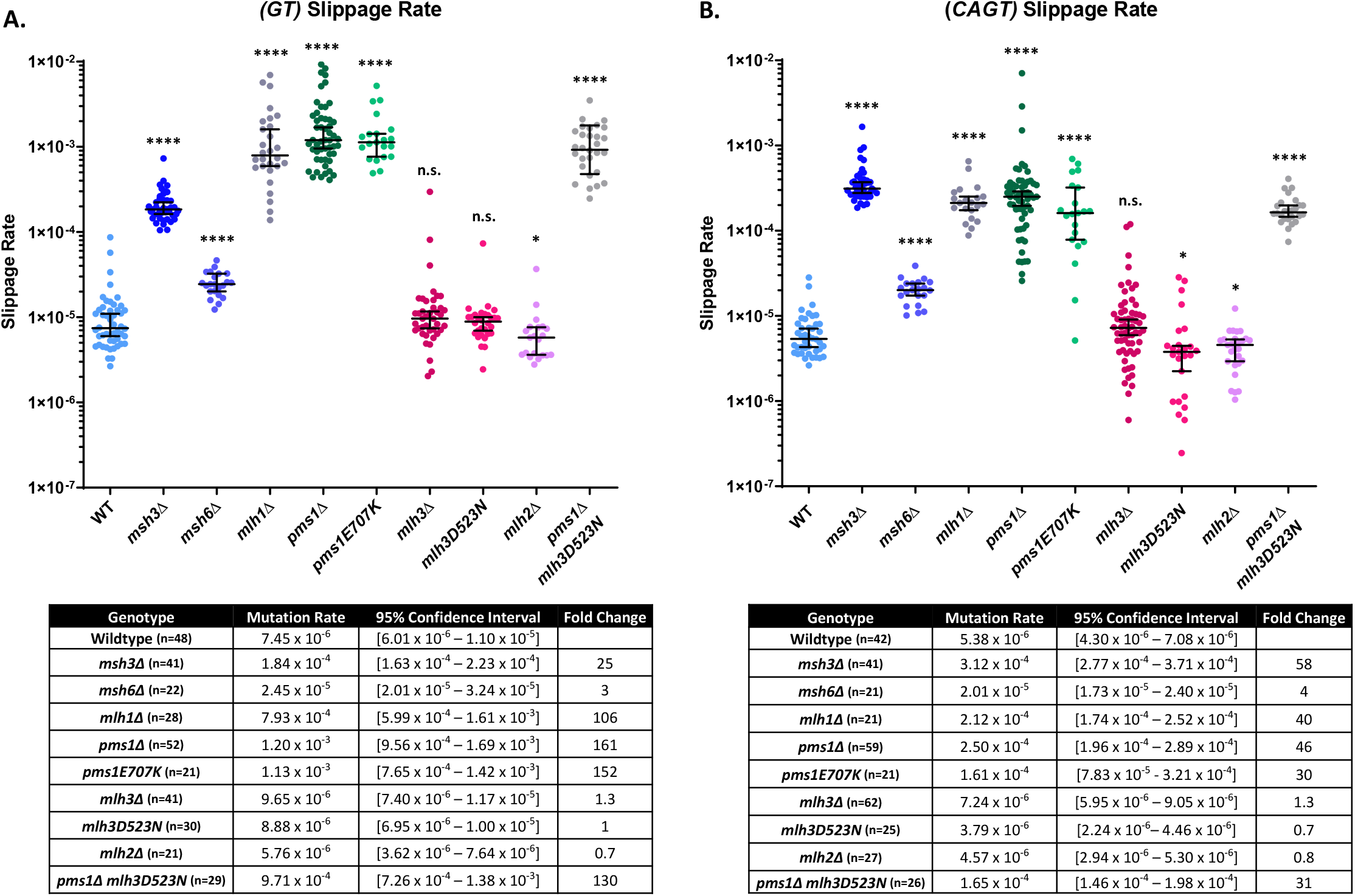
Microsatellite instability of *GT* and *CAGT* tracts. The rate of unrepaired slippage events of *GT* (**A**) and *CAGT* (**B**) were determined in the following genotypes. Median rate and 95% confidence intervals are indicated on the graphs. Tables below include mutation rate, 95% confidence interval, and fold change compared to wild-type. *P*-values were calculated using Mann-Whitney test compared to wild-type: n.s. not significant; \**P* < 0.05; **** *P* < 0.0001

The repeat units are placed in-frame and upstream of the *URA3* gene, allowing production of Ura3, which converts 5-fluoroorotic acid (5-FOA) to the toxic 5-fluorouracil (5-FU), making cells sensitive to 5-FOA (Boeke *et al*. 1987). Unrepaired DNA polymerase-generated slippage events generate IDLs that put *URA3* out of frame, preventing Ura3 production and generating 5-FOA resistance, thus allowing selection of slippage events and calculation of slippage rates in different genetic backgrounds.

As previously observed, *MSH3* played an important role in directing repair of the IDLs generated in the dinucleotide and tetranucleotide tracts. In *msh3Δ*, we observed a ∼25-fold increase in slippage rates of dinucleotide repeats (**Figure 1**). In contrast, loss of *MSH6* resulted in a modest ∼3-fold increase in slippage rate for both repeats (**Figure 1**). We note that the dinucleotide slippage rate in *msh2Δ* was significantly higher in previous studies, with rates ∼350-fold higher than wild-type compared to a ∼50-fold increase in *msh3Δ* and a 2-fold increase in *msh6Δ* (Sia *et al*. 1997; Lee *et al*. 2007). These results are consistent with both Msh2-Msh3 and Msh2-Msh6 complexes contributing to repair of dinucleotide IDLs, with Msh2-Msh3 playing a larger role (Sia *et al*. 1997; Lee *et al*. 2007; Lamb *et al*. 2022; Plavskin *et al*. 2022). In the presence of the tetranucleotide repeat, *msh3Δ* exhibited an ∼58-fold increase in slippage rate, while *msh6Δ* exhibited a modest 4-fold increase. Notably, loss of *MSH2* was epistatic with loss of *MSH3* in the presence of the tetranucleotide repeat in previous studies (Sia *et al*. 1997; Lee *et al*. 2007), indicating that Msh2-Msh3 was almost exclusively responsible for repair of tetranucleotide repeat IDLs.

Loss of *MLH1*, which encodes the common subunit to all three MLH complexes, displayed significantly increased *GT* and *CAGT* slippage rates. In *mlh1Δ* the dinucleotide and tetranucleotide slippage rates increased ∼106-fold and ∼40-fold compared to wild-type, respectively. This is consistent with loss of Msh2-Msh3 and Msh2-Msh6 activity in repair of 2-base IDLs and loss of Msh2-Msh3 in repair of 4-base IDLs, indicating that MSH-mediated IDL repair requires the activity of functional MLH complexes.

In a *pms1Δ* background, which eliminated Mlh1-Pms1 (MutLα), the dinucleotide *GT* slippage was ∼161-fold higher than wild-type, significantly higher than in *msh3Δ* (**Figure 1**). This observation supports earlier work (Sia *et al*. 2001) and is consistent with a loss of both Msh2-Msh3 and Msh2-Msh6 MMR function observed in *msh2Δ* (Sia *et al*. 1997; Lee *et al*. 2007) when Mlh1-Pms1 is eliminated. Notably, we observed an ∼46-fold increase in slippage rate within the *CAGT* repeat over wild-type, somewhat lower than that of *msh3Δ*, which exhibited a 58-fold increase over wild-type. These results are consistent with significant loss of MMR activity on tetranucleotide repeat substrates in the absence of Mlh1-Pms1, but also suggested that another MLH complex contributes specifically to Msh2-Msh3-mediated MMR of tetranucleotide repeats. Overall, our results are consistent with previous results indicating that Mlh1-Pms1 is responsible for the primary MLH function in MMR in repetitive sequences (Prolla *et al*. 1994; Sia *et al*. 2001; Kunkel and Erie 2005; Kunkel and Erie 2015; Pannafino and Alani 2021).

Loss of either Mlh1-Mlh3 (MutLγ) or Mlh1-Mlh2 (MutLβ) resulted in only minor effects on dinucleotide and tetranucleotide slippage rates (**Figure 1**). In *mlh3Δ*, there was no significant increase in slippage rate for either repeat, although there was a minor trend upward in both the *GT* and *CAGT* repeats. In *mlh2Δ*, we observed a small but significant decrease in mutation rate compared to wild-type, a 0.7- and 0.8-fold change in the presence of the *GT* and *CAGT* repeats, respectively. These data indicate that Mlh2 is not only playing a very minor role in MMR, but its presence may have a detrimental effect on repair of these substrates.

### Pms1 endonuclease activity is required for Msh2-Msh3-mediated MMR

In *pms1E707K*, which inactivates Pms1 endonuclease activity (Erdeniz *et al*. 2007; Kadyrov *et al*. 2007), the slippage rate within the *GT* repeat was increased ∼152-fold over wild-type. The slippage rate within the *CAGT* repeat was increased ∼30-fold over wild-type. These results are consistent with the importance of Mlh1-Pms1 endonuclease activity for MMR. We note, however, that, while elevated, the *CAGT* slippage rate was nonetheless lower in *pms1E707K* compared *pms1Δ*. Together these data are consistent with Mlh1-Pms1 endonuclease activity being required for Mlh1-Pms1 function in Msh2-Msh3 and Msh2-Msh6-mediated MMR, and suggests a minor role for Mlh1-Mlh3 endonuclease activity in repair of 4-base IDLs. We also measured slippage rates in *mlh3D523N*, which eliminates the Mlh3 endonuclease activity (Nishant *et al*. 2008; Rogacheva *et al*. 2014). While the *GT* slippage rate was similar to wild-type in *mlh3D523N*, we observed a significant decrease in *CAGT* slippage rate (0.7-fold) compared to wild-type (**Figure 1**). Thus, loss of Mlh1-Mlh3 endonuclease activity slightly improved Msh2-Msh3-mediated MMR compared to wild-type, suggesting some interaction between Mlh1-Pms1 and Mlh1-Mlh3.

To examine the combinatorial loss of both MMR endonucleases, we measured *GT* and *CAGT* slippage rates in *pms1Δ mlh3D523N*. In this background we observed a 130-fold increase in *GT* slippage rate compared to wild-type, similar to the *mlh1Δ* and *pms1Δ* backgrounds. The *CAGT* slippage rate increased 31-fold over wild-type, similar to both *mlh1Δ* and *pms1Δ*. These observations suggest that *pms1Δ* and *mlh3D523N* are epistatic, consistent Mlh1-Mlh3 and its endonuclease activity playing a minor role in MMR and potentially collaborating with Mlh1-Pms1.

### Loss of MLH complexes reduce TNR expansion rates

We performed *in vivo* TNR expansion assays to define the requirement for each of the three MLH complexes in promoting TNR expansions in *S. cerevisiae*. This also allowed a comparison of the relative importance of each MLH complex in Msh2-Msh3-mediated MMR versus TNR expansions. The TNR expansion assay is based on the *URA3*-based reporter system developed by the Lahue lab (MIRET *et al*. 1998; Dixon *et al*. 2004). Briefly, a tract of 25 *CAG* repeats was inserted in the promoter region of the *S. pombe ADH* gene upstream of *URA3*. TNR expansions that result in a tract greater than 29 repeats alter the *URA3* transcriptional start site, preventing *URA3* expression and leading to 5-FOA resistance, allowing selection of expanded TNR tracts. This assay measures the early stages of *CAG* tract expansions, the so-called threshold length tracts. To determine the true expansion rate, the *CAG* tract lengths of a subset of 5-FOA resistant colonies were measured to estimate the proportion of colonies that became 5-FOA resistant independent of tract expansions (Williams *et al*. 2020), potentially due to inactivating mutations within *URA3* or other genes (Casazza *et al*. 2025).

Using this assay, we demonstrated a significant ∼10-fold reduction in *CAG* expansion rate in the absence of *MSH3* (*msh3Δ*) compared to wild-type (**Figure 2**), similar to our previous observations (Kantartzis *et al*. 2012; Williams and Surtees 2015; Casazza *et al*. 2025). The rate of true expansions in wild-type and *msh3Δ* backgrounds were 90% and 66% respectively for *CAG* repeats (Kantartzis *et al*. 2012; Casazza *et al*. 2025) (**Figure 2**). Using this true expansion correction factor for *CAG* expansion rates, we calculated wild-type and *msh3Δ* rates of 2.3 x 10^-6^ and 1.5 x 10^-7^, respectively, an ∼15-fold reduction in the absence of *MSH3*. We note that the corrected rates are estimates because we only quantify a subset of 5-FOA resistant colonies.

**Figure 2.**
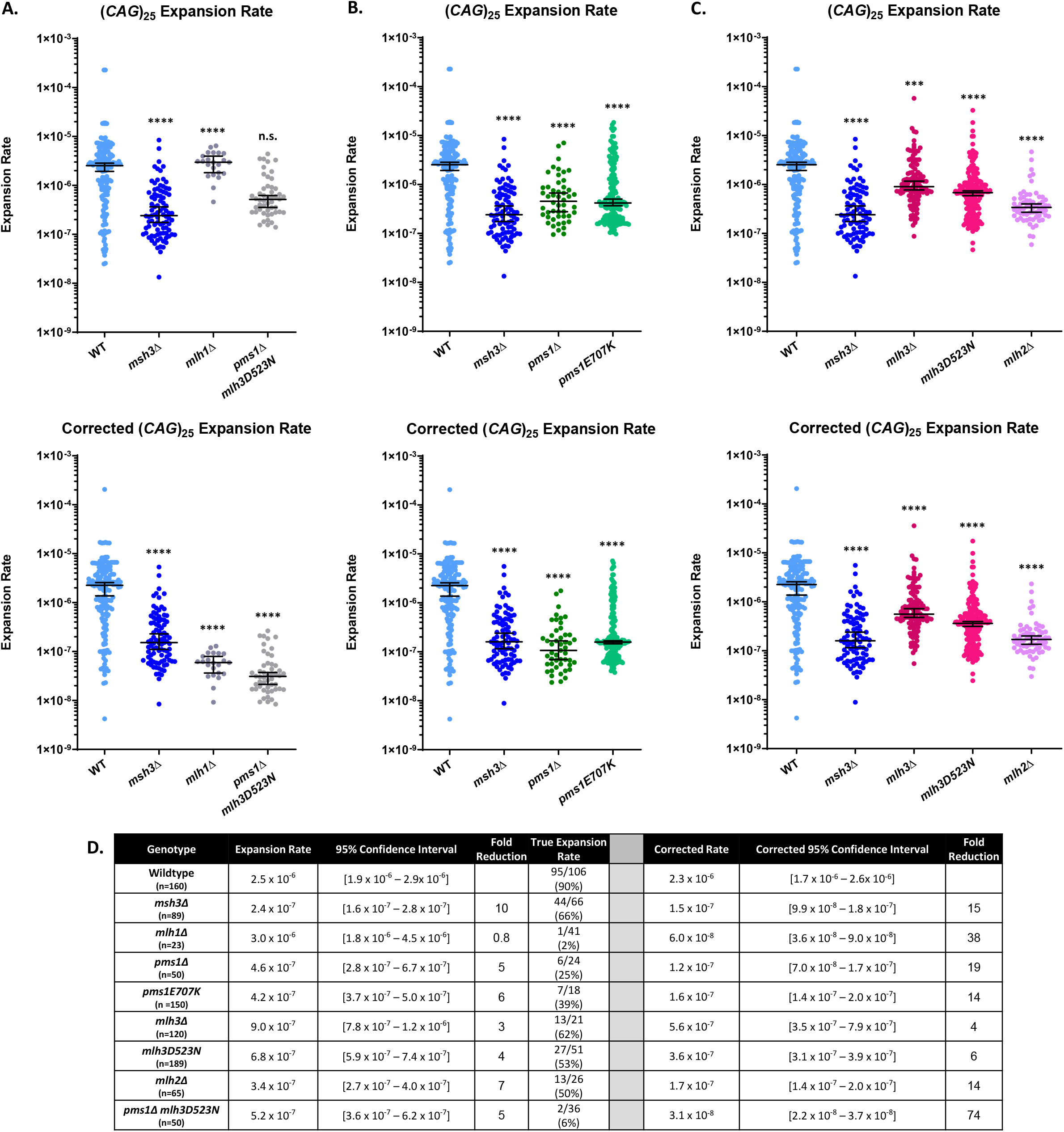
*CAG* expansion rates were determined in the following backgrounds (**A-C**). For each genotype, the uncorrected expansion rate (top panel) and corrected expansion rate (bottom panel) were graphed. The corrected expansion rate was calculated using the true expansion rate (**D**) as a correction factor. Median rate and 95% confidence intervals are indicated on the graphs. Table includes mutation rate, 95% confidence intervals, and fold reduction compared to wild-type for the uncorrected and corrected rates (**D**). *P*-values were calculated using Mann-Whitney test compared to wild-type: n.s. not significant; \*\*\**P* < 0.001; **** *P* < 0.0001

Deletion of *MLH1*, which eliminates all three MLH complexes, resulted in an uncorrected *CAG* expansion rate in *mlh1Δ* was similar to wild-type (**Figure 2A, top panel)**, but with a very low true expansion rate (∼2%). We infer that this low frequency is a result of a significant increase in misincorporation and slippage rates in the absence of *MLH1* (**Figure 1**) (Sia *et al*. 2001; Kunkel and Erie 2005). We previously reported that a significantly elevated mutation rate correlated with a low true expansion rate when characterizing TNR expansion rates in the presence of a chimeric MSH complex (*msh6(3MBD) msh3Δ*) (Casazza *et al*. 2025). Notably, we observed a similarly elevated expansion rate of the scrambled *(C,A,G)*_*25*_ tract in *mlh1Δ* and no true expansions (**Table 3**), consistent with this hypothesis. This is in contrast to the very low expansion rate observed in the wild-type background (**Table 3**) and in *msh3Δ* {kantartzis, 2012}. When we corrected the observed *(CAG)*_*25*_ expansion rate with the true expansion frequency, we observed a very significant decrease in corrected *CAG* expansion rates, ∼38-fold lower than wild-type, which was consistent with loss of Msh2-Msh3-mediated MMR. (**Figure 2A**).

**Table 3:** (*C,A,G*) expansions rates.

| **Genotype** | **Expansion Rate** | **95% Confidence Interval** | **True Expansion Rate** |
| --- | --- | --- | --- |
| **Wildtype (n=89)** | **$<1.0 \times 10^{-8}$** | **-** | **0/40** |
| ***mlh1*Δ (n=30)** | **$1.2 \times 10^{-6}$** | **$8.5 \times 10^{-7} - 2.0 \times 10^{-6}$** | **0/24** |
| ***pms1*Δ (n=40)** | **$8.5 \times 10^{-8}$** | **$1.0 \times 10^{-8} - 1.3 \times 10^{-7}$** | **0/20** |
| ***pms1E707K* (n=30)** | **$1.5 \times 10^{-6}$** | **$9.0 \times 10^{-7} - 2.1 \times 10^{-6}$** | **0/34** |
| ***mlh3*Δ (n=60)** | **$<1.0 \times 10^{-8}$** | **-** | **0/29** |
| ***mlh3D523N* (n=40)** | **$<1.0 \times 10^{-8}$** | **-** | **0/20** |
| ***mlh2*Δ (n=40)** | **$<1.0 \times 10^{-8}$** | **-** | **0/24** |
| ***pms1*Δ *mlh3D523N* (n=77)** | **$1.7 \times 10^{-6}$** | **$1.3 \times 10^{-6} - 2.0 \times 10^{-6}$** | **0/26** |

### Mlh1-Pms1 play a major role in promoting *CAG* expansions

To determine the relative importance of each MLH complex in promoting *CAG* expansions, we determined the *CAG* expansion rate in *pms1Δ*, eliminating Mlh1-Pms1 (MutLα), in *mlh2Δ*, eliminating Mlh1-Mlh2 (MutLβ) or in *mlh3Δ*, eliminating Mlh1-Mlh3 (MutLγ).

*PMS1* has previously been implicated in modifying TNR expansions in several studies (Gomes-Pereira *et al*. 2004; Consortium 2019; Iyer and Pluciennik 2021; Hayward *et al*. 2024). In our system, the *CAG* expansion rate in *pms1Δ* was reduced 5-fold compared to wild-type. As expected, we also observed a low true expansion rate in *pms1Δ* of 25% (**Figure 2B**), consistent with the high mutator phenotype in *pms1Δ* (Sia *et al*. 2001; Romanova and Crouse 2013; Kunkel and Erie 2015) **(Figure 1)**. The corrected *CAG* expansion rate was decreased ∼19-fold, very similar to the rate in *msh3Δ* (**Figure 2B, lower panel**) and higher than the corrected rate for *mlh1Δ* (**Figure 2A**). We also observed a significant reduction in *CAG* expansion rate in *pms1E707K*, the endonuclease-deficient mutation **(Figure 2B)**. The initial measured *CAG* expansion rate was ∼6-fold lower than in wild-type, comparable to *pms1Δ*. The true expansion rate for *pms1E707K* was 39%, intermediate between *pms1Δ* and *msh3Δ* and consistent with an elevated mutation rate (**Figure 1**). The corrected *pm1E707K* expansion rate was ∼14-fold lower than wild-type. Taken together our results indicate that Mlh1-Pms1 is required for *CAG* expansions and makes significant contributions in promoting expansions. Notably, the corrected expansion rate for *mlh1Δ* is significantly lower than that of *pms1Δ*, indicating contributions from one or both of the other MLH complexes. For both *pms1Δ* and *pms1E707K*, we observed elevated apparent scrambled tract expansion rates and no true expansions, again consistent with elevated mutations rates contributing to the uncorrected TNR expansion rates (**Table 3**)

### *MLH2* and *MLH3* make significant contributions to *CAG* expansions

We examined the contribution of Mlh1-Mlh3 (MutLγ) in *CAG* expansions. In *mlh3Δ*, we observed an ∼3-fold decrease in the *CAG* expansion rate compared to wildtype and a corrected rate ∼4-fold lower than wild-type (**Figure 2C**), indicating that Mlh1-Mlh3 contributes to TNR expansions *in viv*o. Our results are consistent with studies implicating *MLH3* as a modifier of *CAG* expansions in both mice and human stem cell models (Pinto *et al*. 2013; Hayward *et al*. 2024). Loss of endonuclease function, with *mlh3D523N*, also significantly reduced the *CAG* expansion rate ∼4-fold, with a corrected rate indicating a ∼6-fold decrease (**Figure 2C**). These results indicate that both Mlh3 and its catalytic activity are required to promote wild-type levels of *CAG* expansions. Notably, this effect, while significant, was smaller than the effect of the loss of *PMS1* (**Figure 2B**).

*MLH2* was notably very important for promoting *CAG* expansions (**Figure 2C**). *mlh2Δ* significantly reduced the expansion rate ∼7-fold lower than wildtype, with a true expansion rate of 50%, resulting in an ∼14-fold reduction compared to wild-type (**Figure 2C**). This decrease is very similar to the decrease observed with *msh3Δ* or *pms1Δ*, indicating a robust requirement for Mlh1-Mlh2 in TNR expansions.

The scrambled tract expansion rates for mutations in *MLH2 and lh3* exhibited very low uncorrected expansion rates, consistent with the low mutator phenotype in these backgrounds (**Table 3**).

### Combinatorial effects of *PMS1* and *MLH3* activities on *CAG* expansions

Our data indicate that both *PMS1* and *MLH3* and their respective endonuclease activities are required for wild-type levels of *CAG* expansions. These results indicate that neither complex can completely compensate for loss of the other and that the endonuclease activity of only one complex is insufficient to promote *CAG* expansions. One possibility is that the two complexes cooperate to promote expansions, as has been suggested for a subset of MMR substrates (Romanova and Crouse 2013). We measured *CAG* expansions rates in *pms1Δ mlh3D523N*, which lacks *PMS1* entirely and is missing the endonuclease activity of *MLH3*, to determine whether these complexes were epistatic with respect to expansion rate. In *pms1Δ mlh3D532N*, we observed an ∼5-fold decrease in *CAG* expansion rate, which was similar to *pms1Δ* alone (**Figure 2A**). Notably, the true expansion rate for *pms1Δ mlh3D523N* was very low (6%) and very similar to *mlh1Δ*. Here, again, the elevated mutator phenotype appears to contribute to the uncorrected TNR expansion rate (**Table 3**). The corrected expansion rate indicates that the double mutant exhibits ∼74-fold decrease in *CAG* expansions, suggesting an additive or potentially synergistic effect of the two mutations.

## Discussion

We have determined the relative contributions of the MLH complexes in both Msh2-Msh3-mediated post-replicative MMR of dinucleotide and tetranucleotides IDLs and *CAG* TNR expansions. Our focus is on the early stages of expansion, i.e. threshold length tracts that transition to pathogenic length tracts. Our yeast model system is uniquely amenable to addressing the mechanistic dynamics at this stage of the expansion process. We observed that, as expected, Mlh1-Pms1 is the primary MLH complex in Msh2-Msh3-mediated MMR, with Mlh1-Mlh2 and Mlh1-Mlh3 making relatively minor contributions. In contrast, all three MLH complexes made substantial contributions promoting *CAG*, tract expansions; loss of any single MLH complex reduced expansion rates significantly, consistent with previous work (Gomes-Pereira *et al*. 2004; Pinto *et al*. 2013; Lahue 2020; Iyer and Pluciennik 2021; Hayward *et al*. 2024). Our data indicated that the MLH complexes are not redundant in *CAG* expansions and that no single MLH complex can fully replace the others in the process.

### Roles of MLH complexes in Msh2-Msh3-mediated repair

To assess MLH complex contributions to Msh2-Msh3-mediated MMR specifically, we focused on slippage events with dinucleotide and tetranucleotide repeats (Sia *et al*. 1997). Msh2-Msh6 plays a less prominent role in repair of both substrates compared with repair of misincorporation events and mononucleotide repeat slippage events and makes only very minor contributions to repair of tetranucleotide repeats (Sia *et al*. 1997; Lee *et al*. 2007; Lamb *et al*. 2022). With both repeats, loss of Mlh1-Pms1 resulted in the most dramatic effect on mutation rates with a >100-fold increase in *GT* slippage events and a >30-fold increase in *CAGT* slippage events (**Figures 1 and 3**), rates similar to loss of *MSH2* (*GT*) or *MSH2* or *MSH3* (*CAGT*) (Sia *et al*. 1997; Sia *et al*. 2001; Lee *et al*. 2007).

**Figure 3:**
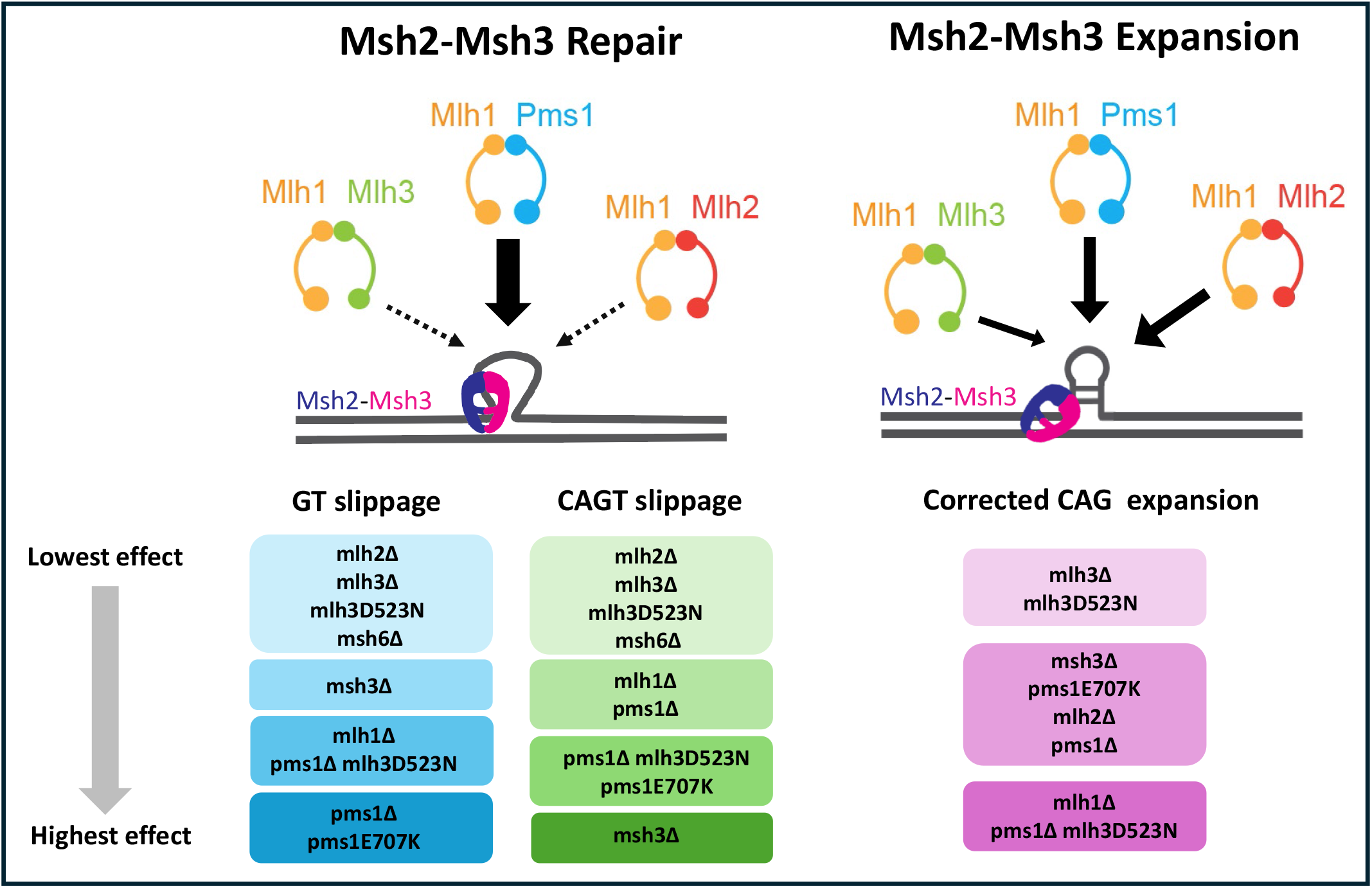
Schematic depicting the contribution each MLH complex makes to Msh2-Msh3 mediated repair and Msh2-Msh3 mediated expansions. The contributions of the MLH complexes vary depending on the pathway (top panel). In appropriate Msh2-Msh3-mediated MMR, Msh2-Msh3 recognizes the IDL structure and recruits Mlh1-Pms1 as the major MLH complex. Mlh1-Mlh3 and Mlh1-Mlh2 make only minor contributions to this pathway. Pathogenic Msh2-Msh3-mediated *CAG* expansions are initiated by Msh2-Msh3 recognizing the *CAG* slipped strand structure in an altered conformation, indicated by the tilted orientation of the complex. All MLH complexes are then recruited and make significant contributions to Msh2-Msh3 mediated expansions, with Mlh1-Pms1 and Mlh1-Mlh2 having more predominant contributions. The bottom panel compares the effect each measured genotype had in Msh2-Msh3 mediated repair versus Msh2-Msh3 mediated expansions as determined experimentally through slippage and expansion assays. Genotypes grouped together in a box denote comparable effects.

Loss of *MLH2* and *MLH3* resulted in more subtle effects that were somewhat repeat-dependent (**Figures 1 and 3**). With both the *GT* and *CAGT* repeats, the slippage rate in *mlh2Δ* was lower than in the wild-type background, suggesting that Mlh1-Mlh2 has a small negative effect on Msh2-Msh3-mediated repair of these substrates. In contrast, Mlh1-Mlh2 was proposed to function as an accessory factor that enhanced Msh2-Msh6-mediated repair (Campbell *et al*. 2014). *MLH2* was shown to be important for repair of large deletion events and 10 nucleotide mononucleotide runs (Harfe *et al*. 2000), indicating Mlh1-Mlh2 is involved in very specific MMR contexts, with a bias toward Msh2-Msh3-mediated events. *MLH3* was shown to be important in repair of −1 and −2 nucleotide deletions, primarily in mononucleotide runs, also consistent with Msh2-Msh3-mediated events (Harfe *et al*. 2000). We observed that loss of *MLH3* resulted in a phenotype indistinguishable from wild-type, but the endonuclease-deficient *mlh3D523N* also decreased the *CAGT* slippage rate below wild-type. This observation suggests a small structural role for Mlh1-Mlh3 in collaborating with Mlh1-Pms1 in MMR, as previously proposed (Romanova and Crouse 2013).

### Roles of MLH complexes in *CAG* expansions

In contrast to Msh2-Msh3-mediated MMR, all three MLH complexes made significant contributions to promoting *CAG* expansions and the endonuclease activity of both Mlh1-Pms1 and Mlhl1-Mlh3 was required for wild-type levels of *CAG* expansions. (**Figures 2 and 3**). These observations are consistent with previous results in mammalian systems that examined the role of individual MLH complexes using much longer, pathogenic repeat tracts (Lahue 2020; Marzec *et al*. 2025). One previous study used a human stem cell model to demonstrate that all three MLH complexes are required for a model of *CAG* expansions (Hayward *et al*. 2024). That work did not, however, characterize the relative contributions of each complex. Our results indicate that loss of either *PMS1* (MutLα) or *MLH2* (MutLβ) decreases expansion rates to an extent similar to that observed with loss of *MSH3*. Loss of *MLH3* (MutLγ) also decreased expansion rates significantly, but the effect was not as substantial. We observed an ∼5-fold decrease with *mlh3Δ* or *mlh3D523N* compared to an ∼15-fold decrease with *msh3Δ, pms1Δ, pms1E707K* or *mlh2Δ*. Furthermore, we found that the effects of *pms1Δ* and *mlh3D523N* were synergistic (**Figures 2 and 3**).

The effect of *mlh2Δ* was particularly notable because Mlh1-Mlh2 does not possess endonuclease activity. This suggests an important structural role for this protein complex. In meiosis, Mlh1-Mlh2 regulates gene conversion tract lengths in a Mer3-dependent manner (Duroc *et al*. 2017). Mlh1-Mlh2 has specificity for D-loop and other branched DNA structures formed during DNA recombination (Duroc *et al*. 2017) and therefore could bind intermediates formed during TNR expansion events. The Mer3-Mlh1-Mlh2 complex also inhibits Pif1 activity, regulating strand displacement synthesis by DNA polymerase δ and RFC-PCNA, suggesting that this complex competes with Pif1 for binding to RFC-PCNA (Vernekar *et al*. 2021). The Mer3-Mlh1-Mlh2 complex also affects RPA filament length during DNA synthesis associated with meiotic recombination (Vernekar *et al*. 2021; Zhai *et al*. 2023). These observations suggest that Mlh1-Mlh2 may coordinate activity of other proteins at the replication fork to promote TNR expansions. Given its minimal involvement in MMR, targeting Mlh1-Mlh2, or Mlh2 specifically, is a potential therapeutic target that would presumably reduce the rate of triplet repeat expansions without impacting normal MMR function (Marzec *et al*. 2025). This would be potentially powerful at a threshold length tract to prevent carriers and afflicted individuals from tract expansions and subsequent disease onset and progression.

### Potential collaboration among MLH complexes

We observed a clear lack of redundancy among the MLH complexes in *CAG* expansions. This is consistent with previous work that has demonstrated the importance of each individual MLH complexes in TNR expansions (Pinto *et al*. 2013; Roy *et al*. 2021; Wang *et al*. 2025). All three MLH complexes were shown to be important in a mouse model of Fragile X *CGG* expansions (Miller *et al*. 2020) and in a human induced pluripotent stem cell model of *CAG* expansions (Hayward *et al*. 2024). These observations suggested the possibility that the three MLH complexes collaborate in this *CAG* expansion process. Previous genetic data supported the hypothesis that Mlh1-Pms1 and Mlh1-Mlh3 cooperate to direct repair of a subset of MMR substrates in yeast (Romanova and Crouse 2013). The role of Mlh1-Mlh2 was not tested in this context.

Mlh1-Pms1 and Mlh1-Mlh3 form filaments along DNA (Hall *et al*. 2001; Manhart *et al*. 2017; Dai *et al*. 2021). Mlh1-Mlh2 has been shown to partially co-localize with Mlh1-Pms1 foci in yeast (Campbell *et al*. 2014), consistent with functional interactions among these MLH complexes that could impact MMR and TNR expansions. Yeast two-hybrid interactions have been demonstrated between Mlh2 and Pms1 (Reyes *et al*. 2020). Mlh3 and Pms1 have been shown to interact through affinity capture (Sanchez *et al*. 2020). Genetic interactions have been reported between *PMS1* and *MLH2* (Campbell *et al*. 2014) and between *PMS1* and *MLH3* (Romanova and Crouse 2013). We suggest that Mlh1-Mlh2 interacts with Mlh1-Pms1, Mlh1-Mlh3 or both complexes to form an MLH heterofilament on the DNA. We propose that these interactions modulate Mlh1-Pms1 and/or Mlh1-Mlh3 endonuclease activity, potentially contributing to the altered strand preference observed *in vitro* in the presence of a TNR slipped strand substrate, resulting in the nicking of the template strand (Pluciennik *et al*. 2013; Kadyrova *et al*. 2020) and potential incorporation of the extrahelical repeats rather than removal.

Importantly, Mlh1-Pms1, Mlh1-Mlh2 and Mlh1-Mlh3 have all also been shown to physically interact with Msh2-Msh3 (Habraken *et al*. 1997; Campbell *et al*. 2014; Rogacheva *et al*. 2014; Al-Sweel *et al*. 2017). This directly connects all three MLH complexes to Msh2-Msh3-mediated pathways. We propose that Msh2-Msh3 promotes two distinct pathways that involve MLH proteins. The first is what we refer to as “appropriate” Msh2-Msh3-mediated repair (**Figure 3, left panel)**. This is post-replicative repair of DNA replication errors to decrease overall mutation rates in the cell. In this pathway, Msh2-Msh3 binds an IDL substrate and recruits primarily Mlh1-Pms1 to initiate repair of the IDL by nicking the nascent strand, followed by excision and resynthesis. Mlh1-Mlh2 and Mlh1-Mlh3 play minor roles in this pathway, which is critical for preventing mutagenesis and potentially carcinogenesis (Prolla *et al*. 1998; Taylor *et al*. 2006; Korhonen *et al*. 2008; Li *et al*. 2020; Deng *et al*. 2021reyes, 2020; Zhang *et al*. 2025).

A “pathogenic” Msh2-Msh3-mediated TNR expansion pathway is initiated by Msh2-Msh3 binding a TNR slipped strand substrate, which adopts an altered conformation relative to Ms2-Msh3 bound to an IDL (**Figure 3, right panel)** (Owen *et al*. 2005; Lang *et al*. 2011; Vernekar *et al*. 2021; Casazza *et al*. 2025) We propose that this altered Msh2-Msh3 conformation leads to the recruitment and formation of all three MLH complexes, possibly a mixed filament along the DNA, that directs MLH endonuclease activity to the template strand, allowing incorporation of extrahelical TNR repeats on the nascent strand. Future work will examine the interactions and functions of each MLH complex in this pathogenic pathway.

## Data Availability Statement

All strains and plasmids are available upon request.

## Funding

This work was supported by American Cancer Society Research Scholar Grant (RSG-142350-01), National Institutes of Health (GM154101) and the National Science Foundation (MCB grant #2325415) to JAS.

## Acknowledgments

We thank members of the Surtees lab for technical assistance and for helpful discussions. We thank Dr. Eric Alani for providing integration plasmids.

## Conflict of Interest

The authors declare they have no conflict of interest.

## References

Al-Sweel, N., V. Raghavan, A. Dutta, V. P. Ajith, L. Di Vietro et al., 2017 mlh3 mutations in baker’s yeast alter meiotic recombination outcomes by increasing noncrossover events genome-wide. PLoS Genet 13: e1006974.

Altmannova, V., M. Firlej, F. Müller, P. Janning, R. Rauleder et al., 2023 Biochemical characterisation of Mer3 helicase interactions and the protection of meiotic recombination intermediates. Nucleic Acids Res 51: 4363–4384.

Boeke, J. D., J. Trueheart, G. Natsoulis and G. R. Fink, 1987 5-Fluoroorotic acid as a selective agent in yeast molecular genetics. Methods Enzymol 154: 164–175.

Campbell, C. S., H. Hombauer, A. Srivatsan, N. Bowen, K. Gries et al., 2014 Mlh2 is an accessory factor for DNA mismatch repair in Saccharomyces cerevisiae. PLoS Genet 10: e1004327.

Casazza, K. M., G. M. Williams, L. Johengen, G. Twoey and J. A. Surtees, 2025 Msh2-Msh3 DNA-binding is not sufficient to promote trinucleotide repeat expansions in Saccharomyces cerevisiae. Genetics 229.

consortium, G.-H., 2015 Identification of Genetic Factors that Modify Clinical Onset of Huntington’s Disease. Cell 162: 516–526.

Consortium, G. M. o. H. s. D., 2019 CAG Repeat Not Polyglutamine Length Determines Timing of Huntington’s Disease Onset. Cell 178: 887-900.e814.

Dai, J., A. Sanchez, C. Adam, L. Ranjha, G. Reginato et al., 2021 Molecular basis of the dual role of the Mlh1-Mlh3 endonuclease in MMR and in meiotic crossover formation. Proceedings of the National Academy of Sciences 118: e2022704118.

De’ Angelis, G. L., L. Bottarelli, C. Azzoni, N. De’ Angelis G. Leandro et al., 2018 Microsatellite instability in colorectal cancer. Acta Biomed 89: 97–101.

Deng, X., M. A. Garcia-Knight, M. M. Khalid, V. Servellita, C. Wang et al., 2021 Transmission, infectivity, and antibody neutralization of an emerging SARS-CoV-2 variant in California carrying a L452R spike protein mutation. medRxiv: 2021.2003.2007.21252647.

Dixon, M. J., S. Bhattacharyya and R. S. Lahue, 2004 Genetic assays for triplet repeat instability in yeast. Methods Mol Biol 277: 29–45.

Drake, J. W., 1991 A constant rate of spontaneous mutation in DNA-based microbes. Proc Natl Acad Sci U S A 88: 7160–7164.

Du, J., E. Campau, E. Soragni, C. Jespersen and J. M. Gottesfeld, 2013 Length-dependent CTG·CAG triplet-repeat expansion in myotonic dystrophy patient-derived induced pluripotent stem cells. Hum Mol Genet 22: 5276–5287.

Duroc, Y., R. Kumar, L. Ranjha, C. Adam, R. Guérois et al., 2017 Concerted action of the MutLβ heterodimer and Mer3 helicase regulates the global extent of meiotic gene conversion. Elife 6.

Erdeniz, N., M. Nguyen, S.M. Deschênes and R. M. Liskay, 2007 Mutations affecting a putative MutLalpha endonuclease motif impact multiple mismatch repair functions. DNA Repair (Amst) 6: 1463–1470.

Flores-Rozas, H., and R. D. Kolodner, 1998 The Saccharomyces cerevisiae MLH3 gene functions in MSH3-dependent suppression of frameshift mutations. Proc Natl Acad Sci U S A 95: 12404–12409.

Foiry, L., L. Dong, C. Savouret, L. Hubert, H. te Riele et al., 2006 Msh3 is a limiting factor in the formation of intergenerational CTG expansions in DM1 transgenic mice. Hum Genet 119: 520–526.

Furman, C. M., R. Elbashir and E. Alani, 2021 Expanded roles for the MutL family of DNA mismatch repair proteins. Yeast 38: 39–53.

Gannon, A. M., A. Frizzell, E. Healy and R. S. Lahue, 2012 MutSβ and histone deacetylase complexes promote expansions of trinucleotide repeats in human cells. Nucleic Acids Res 40: 10324–10333.

Genschel, J., S. J. Littman, J. T. Drummond and P. Modrich, 1998 Isolation of MutSbeta from human cells and comparison of the mismatch repair specificities of MutSbeta and MutSalpha. J Biol Chem 273: 19895–19901.

Gietz, D., A. St Jean, R. A. Woods and R. H. Schiestl, 1992 Improved method for high efficiency transformation of intact yeast cells. Nucleic Acids Res 20: 1425.

Gomes-Pereira, M., M. T. Fortune, L. Ingram, J. P. McAbney and D. G. Monckton, 2004 Pms2 is a genetic enhancer of trinucleotide CAG.CTG repeat somatic mosaicism: implications for the mechanism of triplet repeat expansion. Hum Mol Genet 13: 1815–1825.

Goold, R., J. Hamilton, T. Menneteau, M. Flower, E. L. Bunting et al., 2021 FAN1 controls mismatch repair complex assembly via MLH1 retention to stabilize CAG repeat expansion in Huntington’s disease. Cell Rep 36: 109649.

Gragg, H., B. D. Harfe and S. Jinks-Robertson, 2002 Base composition of mononucleotide runs affects DNA polymerase slippage and removal of frameshift intermediates by mismatch repair in Saccharomyces cerevisiae. Mol Cell Biol 22: 8756–8762.

Gupta, S., M. Gellert and W. Yang, 2011 Mechanism of mismatch recognition revealed by human MutSβ bound to unpaired DNA loops. Nat Struct Mol Biol 19: 72–78.

Habraken, Y., P. Sung, L. Prakash and S. Prakash, 1997 Enhancement of MSH2-MSH3-mediated mismatch recognition by the yeast MLH1-PMS1 complex. Curr Biol 7: 790–793.

Hall, M. C., H. Wang, D. A. Erie and T. A. Kunkel, 2001 High affinity cooperative DNA binding by the yeast Mlh1-Pms1 heterodimer. J Mol Biol 312: 637–647.

Harfe, B. D., and S. Jinks-Robertson, 1999 Removal of frameshift intermediates by mismatch repair proteins in Saccharomyces cerevisiae. Mol Cell Biol 19: 4766–4773.

Harfe, B. D., B. K. Minesinger and S. Jinks-Robertson, 2000 Discrete in vivo roles for the MutL homologs Mlh2p and Mlh3p in the removal of frameshift intermediates in budding yeast. Curr Biol 10: 145–148.

Hayward, B., D. Kumari, S. Santra, C. D. M. van Karnebeek, A. B. P. van Kuilenburg et al., 2024 All three MutL complexes are required for repeat expansion in a human stem cell model of CAG-repeat expansion mediated glutaminase deficiency. Sci Rep 14: 13772.

Higham, C. F., and D. G. Monckton, 2013 Modelling and inference reveal nonlinear length-dependent suppression of somatic instability for small disease associated alleles in myotonic dystrophy type 1 and Huntington disease. J R Soc Interface 10: 20130605.

Higham, C. F., F. Morales, C. A. Cobbold, D. T. Haydon and D. G. Monckton, 2012 High levels of somatic DNA diversity at the myotonic dystrophy type 1 locus are driven by ultra-frequent expansion and contraction mutations. Hum Mol Genet 21: 2450–2463.

Iyer, R. R., and A. Pluciennik, 2021 DNA Mismatch Repair and its Role in Huntington’s Disease. J Huntingtons Dis 10: 75–94.

Jiricny, J., 2006 The multifaceted mismatch-repair system. Nat Rev Mol Cell Biol 7: 335–346.

Kacher, R., F. X. Lejeune, I. David, S. Boluda, G. Coarelli et al., 2024 CAG repeat mosaicism is gene specific in spinocerebellar ataxias. Am J Hum Genet 111: 913–926.

Kadyrov, F. A., S. F. Holmes, M. E. Arana, O. A. Lukianova, M. O’Donnell et al., 2007 Saccharomyces cerevisiae MutLalpha is a mismatch repair endonuclease. J Biol Chem 282: 37181–37190.

Kadyrova, L. Y., V. Gujar, V. Burdett, P. L. Modrich and F. A. Kadyrov, 2020 Human MutLγ, the MLH1-MLH3 heterodimer, is an endonuclease that promotes DNA expansion. Proc Natl Acad Sci U S A 117: 3535–3542.

Kantartzis, A., G. M. Williams, L. Balakrishnan, R. L. Roberts, J. A. Surtees et al., 2012 Msh2-Msh3 interferes with Okazaki fragment processing to promote trinucleotide repeat expansions. Cell Rep 2: 216–222.

Keogh, N., K. Y. Chan, G. M. Li and R. S. Lahue, 2017 MutSβ abundance and Msh3 ATP hydrolysis activity are important drivers of CTG•CAG repeat expansions. Nucleic Acids Res 45: 10068–10078.

Khristich, A. N., J. F. Armenia, R. M. Matera, A. A. Kolchinski and S. M. Mirkin, 2020 Large-scale contractions of Friedreich’s ataxia GAA repeats in yeast occur during DNA replication due to their triplex-forming ability. Proc Natl Acad Sci U S A 117: 1628–1637.

Kirkpatrick, D. T., and T. D. Petes, 1997 Repair of DNA loops involves DNA-mismatch and nucleotide-excision repair proteins. Nature 387: 929–931.

Korhonen, M. K., E. Vuorenmaa and M. Nyström, 2008 The first functional study of MLH3 mutations found in cancer patients. Genes Chromosomes Cancer 47: 803–809.

Kumar, C., G. M. Williams, B. Havens, M. K. Dinicola and J. A. Surtees, 2013 Distinct requirements within the Msh3 nucleotide binding pocket for mismatch and double-strand break repair. J Mol Biol 425: 1881–1898.

Kunkel, T. A., and D. A. Erie, 2005 DNA mismatch repair. Annu Rev Biochem 74: 681–710.

Kunkel, T. A., and D. A. Erie, 2015 Eukaryotic Mismatch Repair in Relation to DNA Replication. Annu Rev Genet 49: 291–313.

Lahue, R. S., 2020 New developments in Huntington’s disease and other triplet repeat diseases: DNA repair turns to the dark side. Neuronal Signal 4: Ns20200010.

Lamb, N. A., J. E. Bard, R. Loll-Krippleber, G. W. Brown and J. A. Surtees, 2022 Complex mutation profiles in mismatch repair and ribonucleotide reductase mutants reveal novel repair substrate specificity of MutS homolog (MSH) complexes. Genetics 221.

Lang, W. H., J. E. Coats, J. Majka, G. L. Hura, Y. Lin et al., 2011 Conformational trapping of mismatch recognition complex MSH2/MSH3 on repair-resistant DNA loops. Proc Natl Acad Sci U S A 108: E837–844.

Lee, J.-M., Z. L. McLean, K. Correia, J. W. Shin, S. Lee et al., 2025 Genetic modifiers of somatic expansion and clinical phenotypes in Huntington’s disease highlight shared and tissue-specific effects. Nature Genetics 57: 1426–1436.

Lee, S. D., J. A. Surtees and E. Alani, 2007 Saccharomyces cerevisiae MSH2-MSH3 and MSH2-MSH6 complexes display distinct requirements for DNA binding domain I in mismatch recognition. J Mol Biol 366: 53–66.

Leeflang, E. P., S. Tavaré, P. Marjoram, C. O. Neal, J. Srinidhi et al., 1999 Analysis of germline mutation spectra at the Huntington’s disease locus supports a mitotic mutation mechanism. Hum Mol Genet 8: 173–183.

Leeflang, E. P., L. Zhang, S. Tavaré, R. Hubert, J. Srinidhi et al., 1995 Single sperm analysis of the trinucleotide repeats in the Huntington’s disease gene: quantification of the mutation frequency spectrum. Hum Mol Genet 4: 1519–1526.

Li, X., Y. Wu, P. Suo, G. Liu, L. Li et al., 2020 Identification of a novel germline frameshift mutation p.D300fs of PMS1 in a patient with hepatocellular carcinoma: A case report and literature review. Medicine (Baltimore) 99: e19076.

Lipkin, S. M., P. B. Moens, V. Wang, M. Lenzi, D. Shanmugarajah et al., 2002 Meiotic arrest and aneuploidy in MLH3-deficient mice. Nat Genet 31: 385–390.

Lipkin, S. M., V. Wang, R. Jacoby, S. Banerjee-Basu, A. D. Baxevanis et al., 2000 MLH3: a DNA mismatch repair gene associated with mammalian microsatellite instability. Nat Genet 24: 27–35.

Mangiarini, L., K. Sathasivam, M. Seller, B. Cozens, A. Harper et al., 1996 Exon 1 of the HD gene with an expanded CAG repeat is sufficient to cause a progressive neurological phenotype in transgenic mice. Cell 87: 493–506.

Manhart, C. M., X. Ni, M. A. White, J. Ortega, J. A. Surtees et al., 2017 The mismatch repair and meiotic recombination endonuclease Mlh1-Mlh3 is activated by polymer formation and can cleave DNA substrates in trans. PLoS Biol 15: e2001164.

Manley, K., J. Pugh and A. Messer, 1999 Instability of the CAG repeat in immortalized fibroblast cell cultures from Huntington’s disease transgenic mice. Brain Res 835: 74–79.

Marsischky, G. T., N. Filosi, M. F. Kane and R. Kolodner, 1996 Redundancy of Saccharomyces cerevisiae MSH3 and MSH6 in MSH2-dependent mismatch repair. Genes Dev 10: 407–420.

Marzec, P., M. Richer and R. S. Lahue, 2025 Therapeutic targeting of mismatch repair proteins in triplet repeat expansion diseases. DNA Repair (Amst) 147: 103817.

Miao, H. K., L. P. Chen, D. P. Cai, W. J. Kong, L. Xiao et al., 2015 MSH3 rs26279 polymorphism increases cancer risk: a meta-analysis. Int J Clin Exp Pathol 8: 11060–11067.

Miller, C. J., G. Y. Kim, X. Zhao and K. Usdin, 2020 All three mammalian MutL complexes are required for repeat expansion in a mouse cell model of the Fragile X-related disorders. PLoS Genet 16: e1008902.

Miret, J. J., L. Pessoa-Brandão and R. S. Lahue, 1998 Orientation-dependent and sequence-specific expansions of CTG/CAG trinucleotide repeats in Saccharomyces cerevisiae. Proc Natl Acad Sci U S A 95: 12438–12443.

Mirkin, S. M., 2007 Expandable DNA repeats and human disease. Nature 447: 932–940.

Nishant, K. T., A. J. Plys and E. Alani, 2008 A mutation in the putative MLH3 endonuclease domain confers a defect in both mismatch repair and meiosis in Saccharomyces cerevisiae. Genetics 179: 747–755.

Owen, B. A., Z. Yang, M. Lai, M. Gajec, J. D. Badger, 2nd et al., 2005 (CAG)(n)-hairpin DNA binds to Msh2-Msh3 and changes properties of mismatch recognition. Nat Struct Mol Biol 12: 663–670.

Pannafino, G., and E. Alani, 2021 Coordinated and Independent Roles for MLH Subunits in DNA Repair. Cells 10.

Paulson, H. L., and K. H. Fischbeck, 1996 Trinucleotide repeats in neurogenetic disorders. Annu Rev Neurosci 19: 79–107.

Pinto, R. M., E. Dragileva, A. Kirby, A. Lloret, E. Lopez et al., 2013 Mismatch repair genes Mlh1 and Mlh3 modify CAG instability in Huntington’s disease mice: genome-wide and candidate approaches. PLoS Genet 9: e1003930.

Plavskin, Y., M. S. de Biase, R. F. Schwarz and M. L. Siegal, 2022 The rate of spontaneous mutations in yeast deficient for MutSβ function. G3 Genes|Genomes|Genetics 13.

Pluciennik, A., V. Burdett, C. Baitinger, R. R. Iyer, K. Shi et al., 2013 Extrahelical (CAG)/(CTG) triplet repeat elements support proliferating cell nuclear antigen loading and MutLα endonuclease activation. Proc Natl Acad Sci U S A 110: 12277–12282.

Porro, A., M. Mohiuddin, C. Zurfluh, V. Spegg, J. Dai et al., 2021 FAN1-MLH1 interaction affects repair of DNA interstrand cross-links and slipped-CAG/CTG repeats. Science Advances 7: eabf7906.

Prolla, T. A., S. M. Baker, A. C. Harris, J. L. Tsao, X. Yao et al., 1998 Tumour susceptibility and spontaneous mutation in mice deficient in Mlh1, Pms1 and Pms2 DNA mismatch repair. Nat Genet 18: 276–279.

Prolla, T. A., Q. Pang, E. Alani, R. D. Kolodner and R. M. Liskay, 1994 MLH1, PMS1, and MSH2 interactions during the initiation of DNA mismatch repair in yeast. Science 265: 1091–1093.

Reyes, G. X., T. T. Schmidt, R. D. Kolodner and H. Hombauer, 2015 New insights into the mechanism of DNA mismatch repair. Chromosoma 124: 443–462.

Reyes, G. X., B. Zhao, T. T. Schmidt, K. Gries, M. Kloor et al., 2020 Identification of MLH2/hPMS1 dominant mutations that prevent DNA mismatch repair function. Commun Biol 3: 751.

Richard, G. F., 2021 The Startling Role of Mismatch Repair in Trinucleotide Repeat Expansions. Cells 10.

Rogacheva, M. V., C. M. Manhart, C. Chen, A. Guarne, J. Surtees et al., 2014 Mlh1-Mlh3, a meiotic crossover and DNA mismatch repair factor, is a Msh2-Msh3-stimulated endonuclease. J Biol Chem 289: 5664–5673.

Romanova, N. V., and G. F. Crouse, 2013 Different roles of eukaryotic MutS and MutL complexes in repair of small insertion and deletion loops in yeast. PLoS Genet 9: e1003920.

Roy, J. C. L., A. Vitalo, M. A. Andrew, E. Mota-Silva, M. Kovalenko et al., 2021 Somatic CAG expansion in Huntington’s disease is dependent on the MLH3 endonuclease domain, which can be excluded via splice redirection. Nucleic Acids Res 49: 3907–3918.

Sanchez, A., C. Adam, F. Rauh, Y. Duroc, L. Ranjha et al., 2020 Exo1 recruits Cdc5 polo kinase to MutLγ to ensure efficient meiotic crossover formation. Proc Natl Acad Sci U S A 117: 30577–30588.

Santos, L. S., S. N. Silva, O. M. Gil, T. C. Ferreira, E. Limbert et al., 2018 Mismatch repair single nucleotide polymorphisms and thyroid cancer susceptibility. Oncol Lett 15: 6715–6726.

Schmidt, M. H. M., and C. E. Pearson, 2016 Disease-associated repeat instability and mismatch repair. DNA Repair (Amst) 38: 117–126.

Senoussi, I., V. Mengoli, A. Cerana, A. Rinaldi, A. Marco et al., 2025 Mechanism of trinucleotide repeat expansion by MutSβ-MutLγ and contraction by FAN1. Nat Commun 16: 9445.

Sia, E. A., M. Dominska, L. Stefanovic and T. D. Petes, 2001 Isolation and characterization of point mutations in mismatch repair genes that destabilize microsatellites in yeast. Mol Cell Biol 21: 8157–8167.

Sia, E. A., R. J. Kokoska, M. Dominska, P. Greenwell and T. D. Petes, 1997 Microsatellite instability in yeast: dependence on repeat unit size and DNA mismatch repair genes. Mol Cell Biol 17: 2851–2858.

Strand, M., M. C. Earley, G. F. Crouse and T. D. Petes, 1995 Mutations in the MSH3 gene preferentially lead to deletions within tracts of simple repetitive DNA in Saccharomyces cerevisiae. Proc Natl Acad Sci U S A 92: 10418–10421.

Taylor, N. P., M. A. Powell, R. K. Gibb, J. S. Rader, P. C. Huettner et al., 2006 MLH3 mutation in endometrial cancer. Cancer Res 66: 7502–7508.

Tomé, S., K. Manley, J. P. Simard, G. W. Clark, M. M. Slean et al., 2013 MSH3 polymorphisms and protein levels affect CAG repeat instability in Huntington’s disease mice. PLoS Genet 9: e1003280.

Vernekar, D. V., G. Reginato, C. Adam, L. Ranjha, F. Dingli et al., 2021 The Pif1 helicase is actively inhibited during meiotic recombination which restrains gene conversion tract length. Nucleic Acids Res 49: 4522–4533.

Wang, N., S. Zhang, P. Langfelder, L. Ramanathan, F. Gao et al., 2025 Distinct mismatch-repair complex genes set neuronal CAG-repeat expansion rate to drive selective pathogenesis in HD mice. Cell 188: 1524-1544.e1522.

Warren, J. J., T. J. Pohlhaus, A. Changela, R. R. Iyer, P. L. Modrich et al., 2007 Structure of the human MutSalpha DNA lesion recognition complex. Mol Cell 26: 579–592.

Williams, G. M., A. K. Petrides, L. Balakrishnan and J. A. Surtees, 2020 Tracking Expansions of Stable and Threshold Length Trinucleotide Repeat Tracts In Vivo and In Vitro Using Saccharomyces cerevisiae. Methods Mol Biol 2056: 25–68.

Williams, G. M., and J. A. Surtees, 2015 MSH3 Promotes Dynamic Behavior of Trinucleotide Repeat Tracts In Vivo. Genetics 200: 737–754.

Williams, G. M., and J. A. Surtees, 2018 Measuring Dynamic Behavior of Trinucleotide Repeat Tracts In Vivo in Saccharomyces cerevisiae. Methods Mol Biol 1672: 439–470.

Winston, F., C. Dollard and S. L. Ricupero-Hovasse, 1995 Construction of a set of convenient Saccharomyces cerevisiae strains that are isogenic to S288C. Yeast 11: 53–55.

Yamada, M., T. Sato, S. Tsuji and H. Takahashi, 2008 CAG repeat disorder models and human neuropathology: similarities and differences. Acta Neuropathol 115: 71–86.

Zhai, B., S. Zhang, B. Li, J. Zhang, X. Yang et al., 2023 Dna2 removes toxic ssDNA-RPA filaments generated from meiotic recombination-associated DNA synthesis. Nucleic Acids Res 51: 7914–7935.

Zhang, C., J. Ye, Q. Li, J. Zhan, D. Yang et al., 2025 Follow-up of hereditary endometrial carcinoma caused by MLH3 gene mutation: a case report. Frontiers in Oncology Volume 15 -2025.

